# Impact of extrinsic incubation temperature on natural selection during Zika virus infection of *Aedes aegypti*

**DOI:** 10.1101/2021.03.02.433538

**Authors:** Reyes A. Murrieta, Selene Garcia-Luna, Deedra J. Murrieta, Gareth Halladay, Michael C. Young, Joseph R. Fauver, Alex Gendernalik, James Weger-Lucarelli, Claudia Rückert, Gregory D. Ebel

**Author notes:** Corresponding author: Gregory D. Ebel.

## Abstract

Arthropod-borne viruses (arboviruses) require replication across a wide range of temperatures to perpetuate. While vertebrate hosts tend to maintain temperatures of approximately 37°C - 40°C, arthropods are subject to ambient temperatures which can have a daily fluctuation of > 10°C. Temperatures impact vector competence, extrinsic incubation period, and mosquito survival unimodally, with optimum occurring at some intermediate temperature. In addition, the mean and range of daily temperature fluctuations influence arbovirus perpetuation and vector competence. The impact of temperature on arbovirus genetic diversity during systemic mosquito infection, however, is poorly understood. Therefore, we determined how constant extrinsic incubation temperatures of 25°C, 28°C, 32°C, and 35°C control Zika virus (ZIKV) vector competence and population dynamics within *Aedes aegypti* and *Aedes albopictus* mosquitoes. We also examined diurnally fluctuating temperatures which more faithfully mimic field conditions in the tropics. We found that vector competence varied in a unimodal manner for constant temperatures peaking between 28°C and 32°C for both *Aedes* species. Transmission peaked at 10 days post-infection for *Aedes aegypti* and 14 days for *Aedes albopictus.* The effect of diurnal temperature was distinct and could not have been predicted from constant temperature-derived data. Using RNA-seq to characterize ZIKV population structure, we identified that temperature alters the selective environment in unexpected ways. During mosquito infection, constant temperatures more often elicited positive selection whereas diurnal temperatures led to strong purifying selection in both *Aedes* species. These findings demonstrate that temperature has multiple impacts on ZIKV biology within mosquitoes, including major effects on the selective environment within mosquitoes.

**Author Summary:** Arthropod-borne viruses (arboviruses) have emerged in recent decades due to complex factors that include increases in international travel and trade, the breakdown of public health infrastructure, land use changes, and many other factors. Climate change also has the potential to shift the geographical ranges of arthropod vectors, consequently increasing the global risk of arbovirus infection. Changing temperatures may also alter the virus-host interaction, ultimately resulting in the emergence of new viruses and virus genotypes in new areas. Therefore, we sought to characterize how temperature (both constant and fluctuating) alters the ability of *Aedes aegypti* and *Aedes albopictus* to transmit Zika virus, and how it influences virus populations within mosquitoes. We found that intermediate temperatures maximize virus transmission compared to more extreme and fluctuating temperatures. Constant temperatures increased positive selection on virus genomes, while fluctuating temperatures strengthened purifying selection. Our studies provide evidence that in addition to altering VC, temperature significantly influences the selective environment within mosquitoes.

## Introduction

Arthropod-borne viruses (arboviruses) such as Zika virus (ZIKV, Flaviviridae, *Flavivirus*) are mainly RNA viruses that are transmitted by arthropod vectors among vertebrate hosts [1]. Thus, arboviruses are required to alternately replicate in hosts with drastically different body temperatures. This affects transmission dynamics, replication rates, and population structure. While replication in vertebrates generally occurs within 2-3 degrees of 38°C [2], infection in mosquitoes may occur at a much wider range of temperatures: Mosquito vectors are distributed throughout tropical and temperate climates and the geographical range of important species is increasing [3]. Climate variations such as heat waves, cold snaps, or daily temperature fluctuations change the host environment within which arboviruses replicate and are transmitted. Fluctuations in the temperature of the host environment are central to arbovirus biology [4] and virus-host interaction [5–7].

The impact of temperature on vector competence (VC), i.e. the ability of a mosquito to acquire, maintain, and transmit a pathogen, is well described. Temperature increases impact VC in a unimodal manner (having one clear peak), with extreme low (16°C) and high (38°C) temperatures having low VC while median temperatures (28°C −32°C) have higher VC [8, 9]. The extrinsic incubation temperature (EIT) also influences viral replication and dissemination within vectors [10–15], altering the extrinsic incubation period, i.e. the number of days between acquisition of an infection and infectiousness to a new host [5, 16]. Most studies examining the effects of temperature on VC have used single, constant temperatures to represent optimal conditions for mosquito colony survival [17–20]. However, diurnal temperature fluctuations more accurately model the environmental conditions encountered by mosquitoes in the field [21–24]. Although temperature clearly exerts a strong selective pressure on RNA viruses [25, 26], little is known about how it may influence the composition of arbovirus populations during mosquito infection. Thus, while temperature clearly effects arbovirus transmission and epidemiology, its impact on arbovirus evolution remains unclear.

RNA viruses like ZIKV have the capacity to evolve rapidly in response to changing environments. This is due, in part, to short generation times and error-prone replication [27, 28]. As a result, arboviruses, including ZIKV, exist within hosts as large populations of mixed haplotypes, which is critical to their perpetuation in nature [29–33]. While there have been numerous studies assessing ZIKV VC and viral ecology and some efforts focusing on the use of environmental data to predict virus spread, there is limited knowledge as to how environmental factors such as temperature impact the selective environments and mutational diversity of arboviruses within mosquitoes. Accordingly, we sought to determine whether ZIKV mutational diversity is altered by EIT during systemic infection of *Ae. aegypti* and *Aedes albopictus (Ae. albopictus*) vectors. We exposed mosquitoes to a Puerto Rican isolate of ZIKV and held them at constant temperatures of 25°C, 28°C, 32°C, 35°C; and a diurnal fluctuation from 25°C to 35°C. We then assessed VC and measured virus mutational diversity in different tissue compartments of each mosquito using next-generation sequencing (NGS). Our results suggest that the selective environment within mosquitoes is significantly modified by temperature, and that temperature fluctuations exert unique constraints upon arbovirus sequences.

## Results

To assess how extrinsic incubation temperature affects vector competence for ZIKV we exposed *Ae. aegypti* and *Ae. albopictus* to ZIKV (n=72-108), held them at 25°C, 28°C, 32°C, 35°C, and alternating diurnal temperature that fluctuated between 25°C-35°C. Infection rates were high in all mosquitoes except those held at 35°C (Fig 1). In both *Aedes* species, moderate temperatures (28°C and 32°C) significantly (p < 0.05, Two-tailed Fisher’s exact test) increased dissemination and transmission at 7 days post-infection (Fig 1A & 1C). The difference in dissemination was most notable in *Ae. albopictus,* which was ~30% at 28°C and ~80% at 32°C (Fig 1A). Our diurnal temperature group did not fit with the expected unimodal distribution given the mean daily temperature in this group (30°C). Instead, infection in this temperature group was lower, and most closely resemble infection rates of the 25°C and 28°C temperatures or 32°C and 35°C temperatures. Mosquitoes experiencing diurnal temperatures also had significantly lower dissemination and transmission (p < 0.05, Two-tailed Fisher’s exact test) compared to the standard laboratory colony temperature of 28°C, which is used for most VC studies. [diurnal temperature depressed aedes vector competence compared to the optimal, mean temperature.]

**Fig 1.**
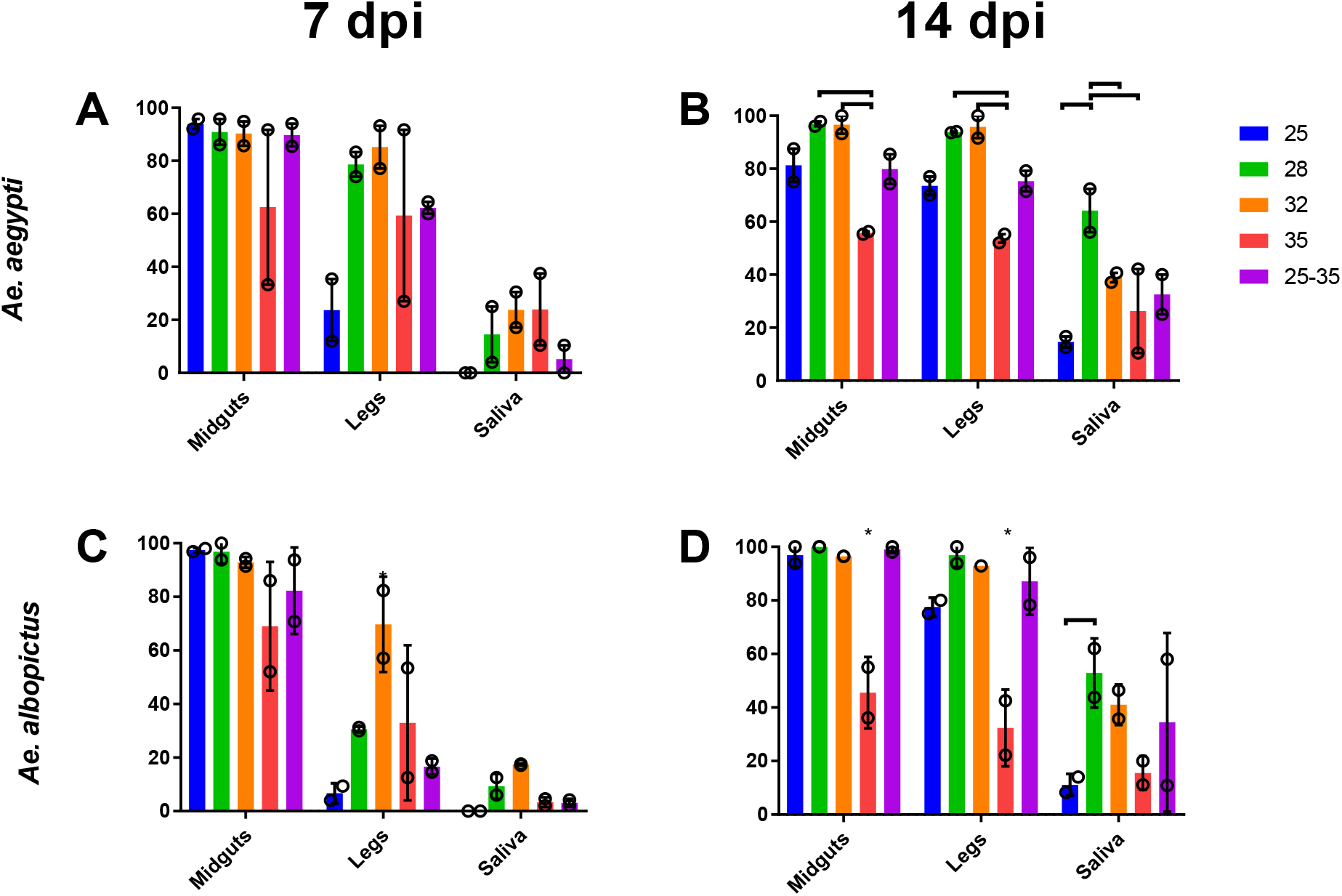
Extrinsic incubation temperature alters ZIKV transmission efficiency in *Aedes* mosquitoes. Percent of *Ae. aegypti* (A & B) and *Ae. albopictus* (C & D) with ZIKV in midgut, legs, and saliva at 7 (A & C) and 14 (B&D) days post feeding. The bar represents the mean and the open circles represent the value of each experiment with SEM shown with error bars. Brackets above vertical bars indicate statistical significance between indicated groups, asterisks indicate significance from the entire species-tissue group (p < 0.05, Two-tailed Fisher’s exact test).

Consensus-level changes were rarely observed in ZIKV in these experiments. We therefore assessed the effect of temperature and mosquito species on ZIKV mutational diversity at the intrahost level by sequencing virus collected from 3 biological replicates of tissues (midgut, legs, and saliva) harvested from *Ae. aegypti* and *Ae. albopictus* at 14 days post exposure. Nucleotide diversity across the coding sequence was lowest in mosquitoes held at 25°C and highest in mosquitoes held at 35°C or under a diurnally fluctuating temperature regime. Minimal differences were observed when comparing *Ae. aegypti* to *Ae. albopictus* (Fig 2A). *d_N_/d_S_* was estimated across the coding sequence to assess selection acting upon viral genomes. In both *Aedes* species, *d_N_/d_S_* was significantly lower in ZIKV from diurnal-exposed mosquitoes compared to those held at constant temperatures, and solely virus from these mosquitoes had *d_N_/d_S_* much lower than 1 (Fig 2B). Similar levels of richness, complexity, and divergence were observed in all mosquitoes and EIT groups (Not Shown).

**Fig 2.**
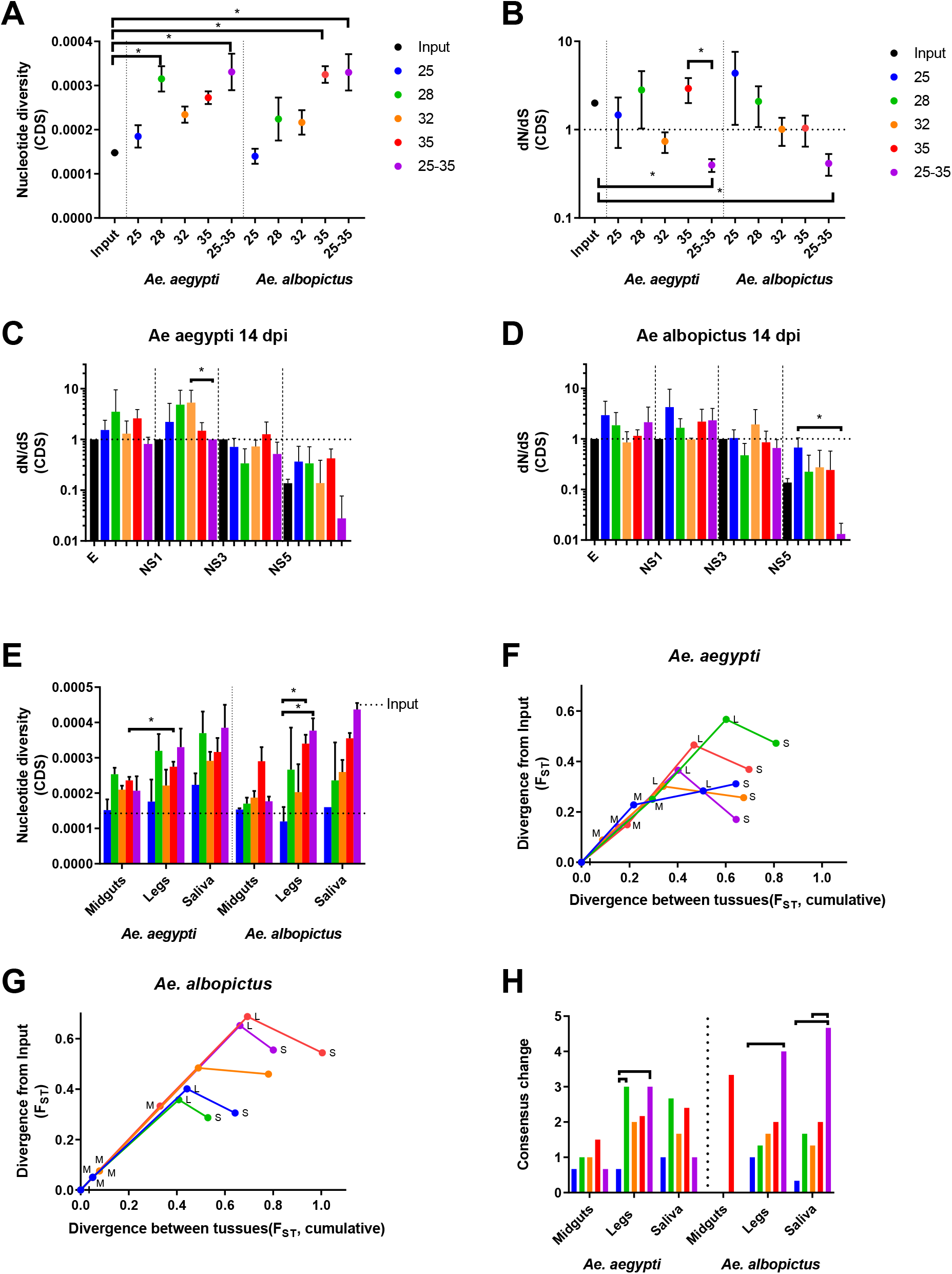
Temperature alters virus diversification, selection and divergence during mosquito infection. ZIKV population diversity at varying temperatures was determined using measures of nucleotide diversity (A) and natural selection (*d_N_/d_S_*) (B). *d_N_/d_S_* was also examined by virus coding region for virus that replicated in *Ae. aegypti* (C), and *Ae. albopictus* (D). Results for E, NS1, NS3, and NS5 protein coding regions are shown. *d_N_/d_S_* was near 1 for C, prM NS2A, NS2B, NS4A, NS4B (not shown. Nucleotide diversity (E) and divergence (F-G) were determined for ZIKV in different mosquito compartments. Divergence from input population (y-axis) and cumulative divergence between tissues (x-axis) (F-G) is presented. Midguts (M), legs (L), Saliva (S). Consensus change counts also are presented (H) by temperature and mosquito species., and majority variants accumulated (H), as markers of population diversity. Significance was tested using the Kruskal-Wallis test with Dunn’s correction (A-D, H, * p < 0.05) or 2-way ANOVA with Tukey’s (E, p-value < 0.05). Figures present the mean and SEM (A-B, E) or 95% CI (C-D).

To assess coding region-specific signatures of selection, we analyzed *d_N_/d_S_* for each viral protein coding sequence independently. In both mosquito species exposed to diurnal temperatures, *d_N_/d_S_* was much less than 0.1 only within the NS5 coding sequence (Fig 2C-D mean *d_N_/d_S_* 0.027 in *Ae. aegypti* and 0.0132 in *Ae. albopictus).* E and NS1 coding sequences had *d_N_/d_S_* greater than 1 when temperatures were constant, ranging from a mean *d_N_/d_S_* of 1.298 at 32°C in E to 5.325 at 32°C in NS1 in *Ae. aegypti. Ae. albopictus* had a mean low *dn/d_S_* of 1.154 at 35°C in E and a high of 4.267 at 25°C in NS1, the exception being 32°C in *Ae. albopictus* where *d_N_/d_S_* was 0.0853 (E) and 0.9804 (NS1) respectively (Fig 2C-D).

Since arboviruses encounter multiple replication environments and barriers during systemic mosquito infection, we next assessed intrahost population diversity in the midguts, legs, and saliva of mosquitoes held at varying temperatures (Fig 2E-H). In all tissue compartments of both species tested, nucleotide diversity tended to increase with increasing temperature (Fig 2E), with diurnally-exposed mosquitoes having lower diversity during midgut infection but increased genetic diversity during systemic infection that resulted in some of the highest levels of nucleotide diversity in leg and saliva-associated virus (Fig 2E). Analysis of the fixation index (F_ST_), during systemic infection revealed temperature- and species-specific patterns of divergence from the input population (Fig 2F-G). Generally, ZIKV diverged more in the midguts of *Ae. aegypti* than *Ae. albopictus.* The 28°C EIT group diverged more than any other EIT group in *Ae. aegypti* (Fig 2F). Whereas in *Ae. albopictus,* exposure to higher temperatures of 32°C and 35°C promote divergence. In both species, divergence was greatest when the population disseminated from the midgut to the legs and decreased from legs to saliva. These data provide evidence that divergence from the founding population was increased in the midgut and legs of both species and reduced as virus moved from legs to saliva, potentially due to stochastic reductions caused by bottlenecks and/or purifying selection.

Patterns of ZIKV population diversity during systemic infection are temperature dependent. This also is reflected in the number of changes to the consensus virus sequence (Fig 2H). Increasing temperature tended to increase the number of consensus changes, with 28°C somewhat of an outlier. Significantly more consensus changes were observed in *Ae. albopictus* for diurnal temperatures than any constant temperature group (Fig 2H).

Since ZIKV population diversity is influenced by the tissues of origin, as well as the constant and diurnal EIT, we assessed *d_N_/d_S_* during systemic infection for each EIT group across the entire CDS, and for the structural and nonstructural regions independently (Fig 3). Our input population had a *d_N_/d_S_* ratio of 1.75 for the CDS, 3.11 for the structural region, and 0.95 for the non-structural regions. This indicates that the structural regions of our input population were under positive selection (*d_N_/d_S_* greater than1) during its propagation and preparation, whereas the non-structural regions were not. Interestingly, when ZIKV was exposed to diurnal fluctuating temperatures, it was under strong purifying selection (*d_N_/d_S_* less than 1) in both *Aedes* species (Fig 3E & 3J), whereas all constant temperatures caused ZIKV *d_N_/d_S_* generally near or above 1 (Fig 3). In the saliva, 25°C and 32°C EIT groups had a *d_N_/d_S_* that neared 1, decreasing from the input in both *Aedes* species (Fig 3A, 3C, 3F, 3H). Conversely, 28°C and 35°C EIT groups maintained or increased *d_N_/d_S_* when compared to input (Fig 3B, 3D, 3G, 3I).

**Fig 3.**
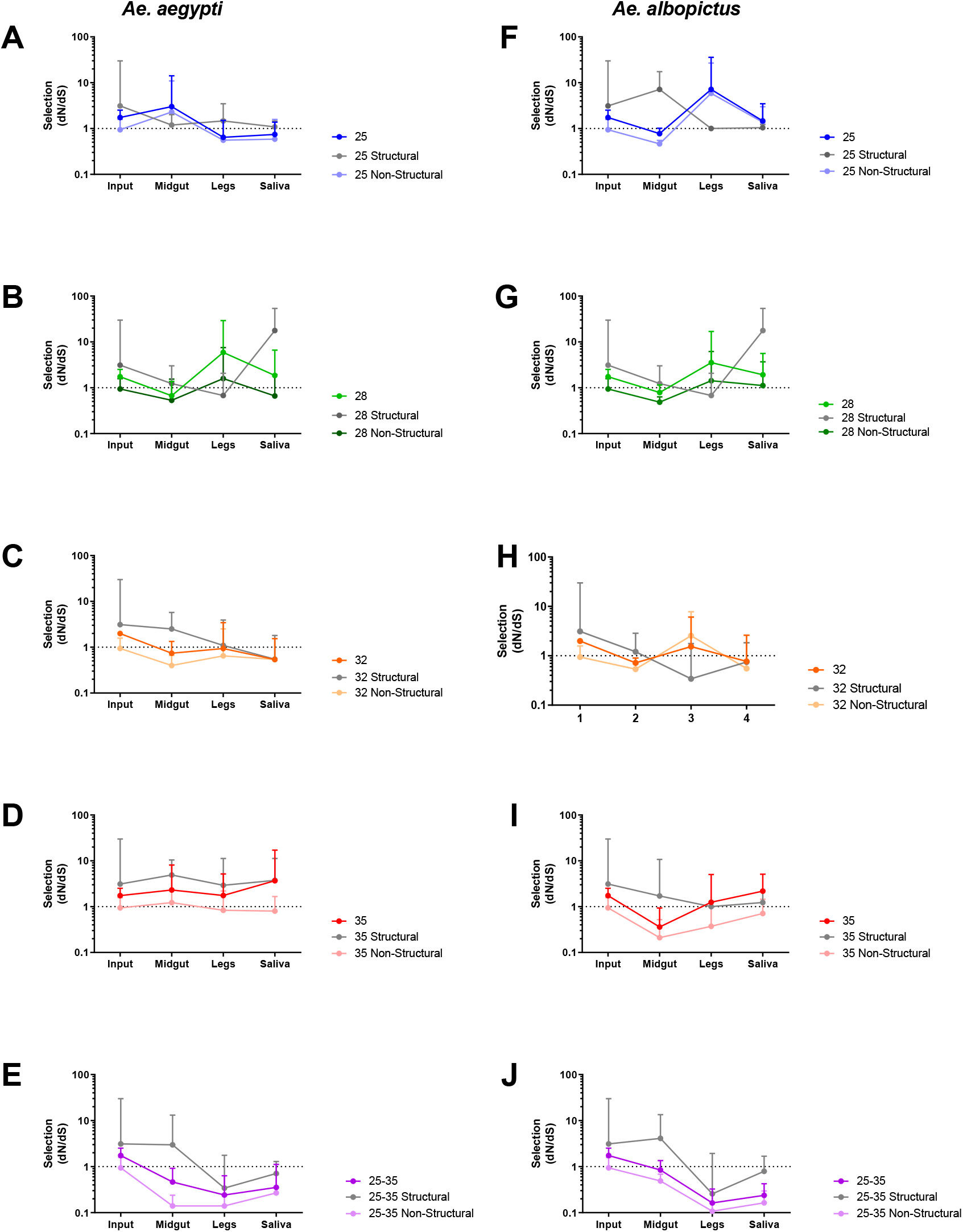
Fluctuating diurnal temperatures impose purifying selection on ZIKV during systemic mosquito infection. *d_N_/d_S_* (mean with 95% CI) for ZIKV CDS (closed circles), structural sequence (Boxes). And Non-structural sequence (open circles), at indicated temperatures, including diurnal fluctuating temperatures, in *Ae. aegypti* (A-E) and *Ae. albopictus* (F-J).

Although relatively few consensus level changes occurred in ZIKV after replication in mosquitoes, we identified a handful of changes to the ZIKV genome that occurred independently in several mosquito tissues (Table 1). Of these consensus changes, 3 nonsynonymous and 1 synonymous SNVs were found in both *Ae. aegypti* and *Ae. albopictus* (L330V E, W98G NS1, M220T NS1, and G83 NS5) samples. The remaining 4 consensus changes were comprised of 1 non-synonymous mutation (T315I E) unique to *Ae. aegypti* and 3 mutations unique to *Ae. albopictus:* 1 non-synonymous mutation and 2 synonymous mutations (K146E NS1, I94 NS2A and F682 NS5). All were present as minority variants in the input virus population, and several have been documented in ZIKV genomic epidemiologic studies. (Table 1). These consensus changes tended to rise or fall in frequency in a species- and temperature-dependent manner (Figure 4). In general, high and/or diurnal temperature tended to drive variants to higher frequency in the population, while at the low temperature (25°C) variants tended to remain closer to their input frequency. The frequency of the synonymous variant G83 in NS5 fluctuated in frequency similarly to M220T, suggesting linkage on the viral genome.

**Table 1.** Multiple ZIKV variants found in across biological samples and temperatures go to consensus. ^a^ Variant frequency found in the stock input ZIKV population. ^b^ The percent sequence identity observed in nature when aligned to 150 complete ZIKV genomes. ^c^ Extrinsic incubation temperatures at which each variant was observed, ^d^ and tissues that each variant was observed. Black= both species, Blue= Ae. aegypti only, Red= Ae. albopictus only. M, midguts; L, legs; S, saliva.

| Species | Coding Sequence | AA | Input Freq<sup>a</sup> | Freq found<sup>b</sup><br>in Nature | Temperature Obs.<sup>c</sup> | Tissue<sup>d</sup><br>Obs. |
| --- | --- | --- | --- | --- | --- | --- |
| *Aedes spp.* | E | L330V | 0.35 | 100% | 28, 32, 35, 25-35 | M, L, S |
|  | NS1 | W98G | 0.30 | 0% | 28, 32, 35, 25-35 | M, L, S |
|  | NS1 | M220T | 0.14 | 0% | 25, 28, 32, 35, 25-35 | M, L, S |
|  | NS5 | G83 | 0.13 | 0% | 25, 28, 32, 35, 25-35 | M, L, S |
| *Ae. aegypti* | E | T315I | 0.05 | 0% | 28, 35 | M, L, S |
| *Ae. albopictus* | NS1 | K146E | 0.01 | 2% | 25, 28, 32, 25-35 | L, S |
|  | NS2A | I94 | 0.03 | 0.7% | 35, 25-35 | M, L, S |
|  | NS5 | F682 | 0.03 | 0% | 35, 25-35 | M, L, S |

**Fig 4.**
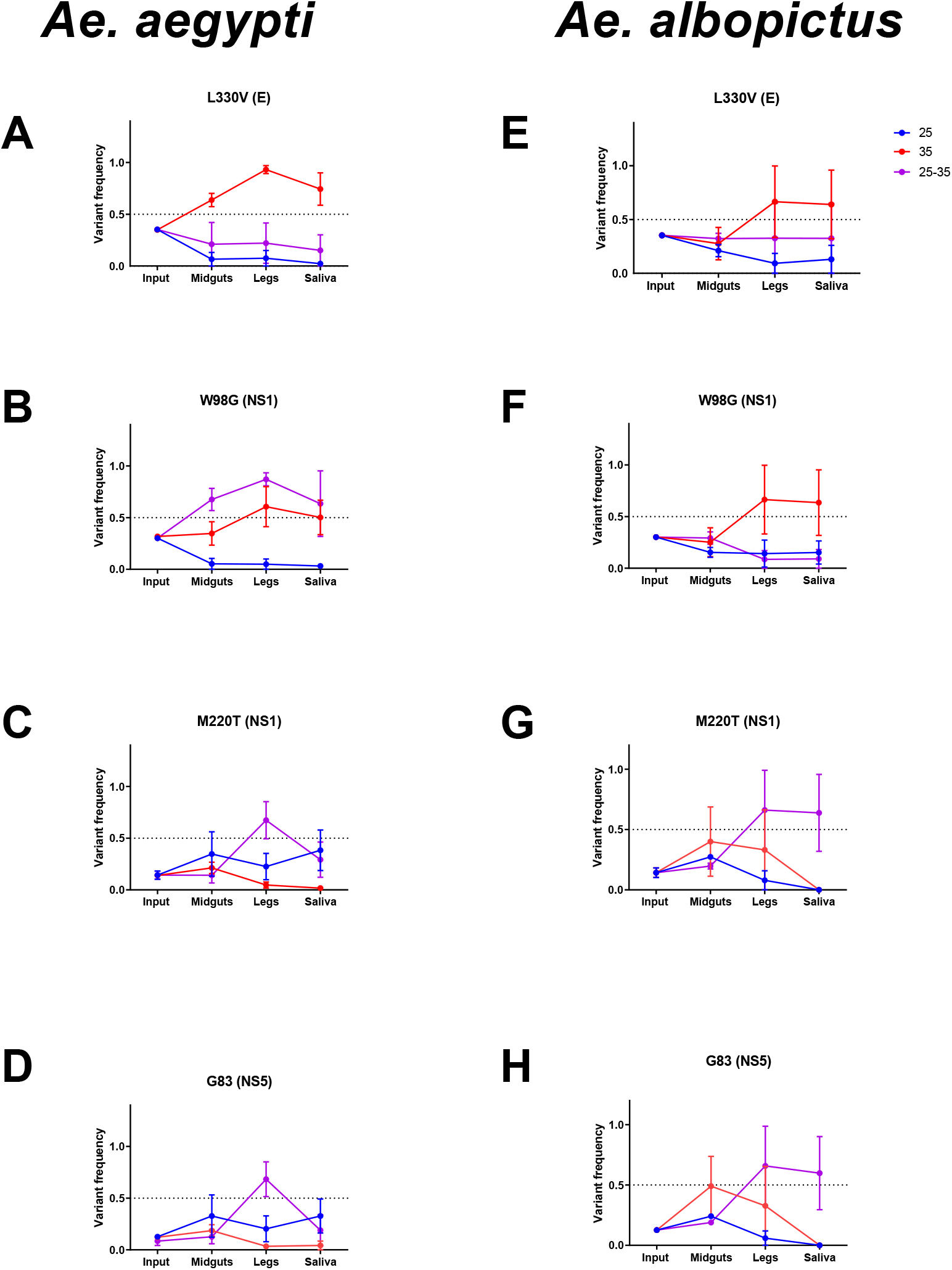
EIT and species control ZIKV variant frequency during systemic infection. Frequencies of L330V E (A & E), W98G NS1 (B & F), M220T NS1 (C & G), and G83 NS5 (D & H) in input, midgut, legs and saliva shown at 25, 35 and diurnal 25-35C in *Ae. aegypti* (A-D) and *Ae. albopictus* (E-H). Mean and SEM of competitions from three biological replicates shown. We then competed engineered ZIKV mutants containing the eight consensus-changing mutations that arose during systemic infection (Table 1) in mosquitoes under low (25°C) and high (35°C) EITs (Figure 5) to assess whether the fitness of these variants may be temperature-dependent. Control competitions with marked and unmarked clones of the PRVABC59 virus were unremarkable, with no significant changes in test to reference virus detected at either EIT.

**Fig 5.**
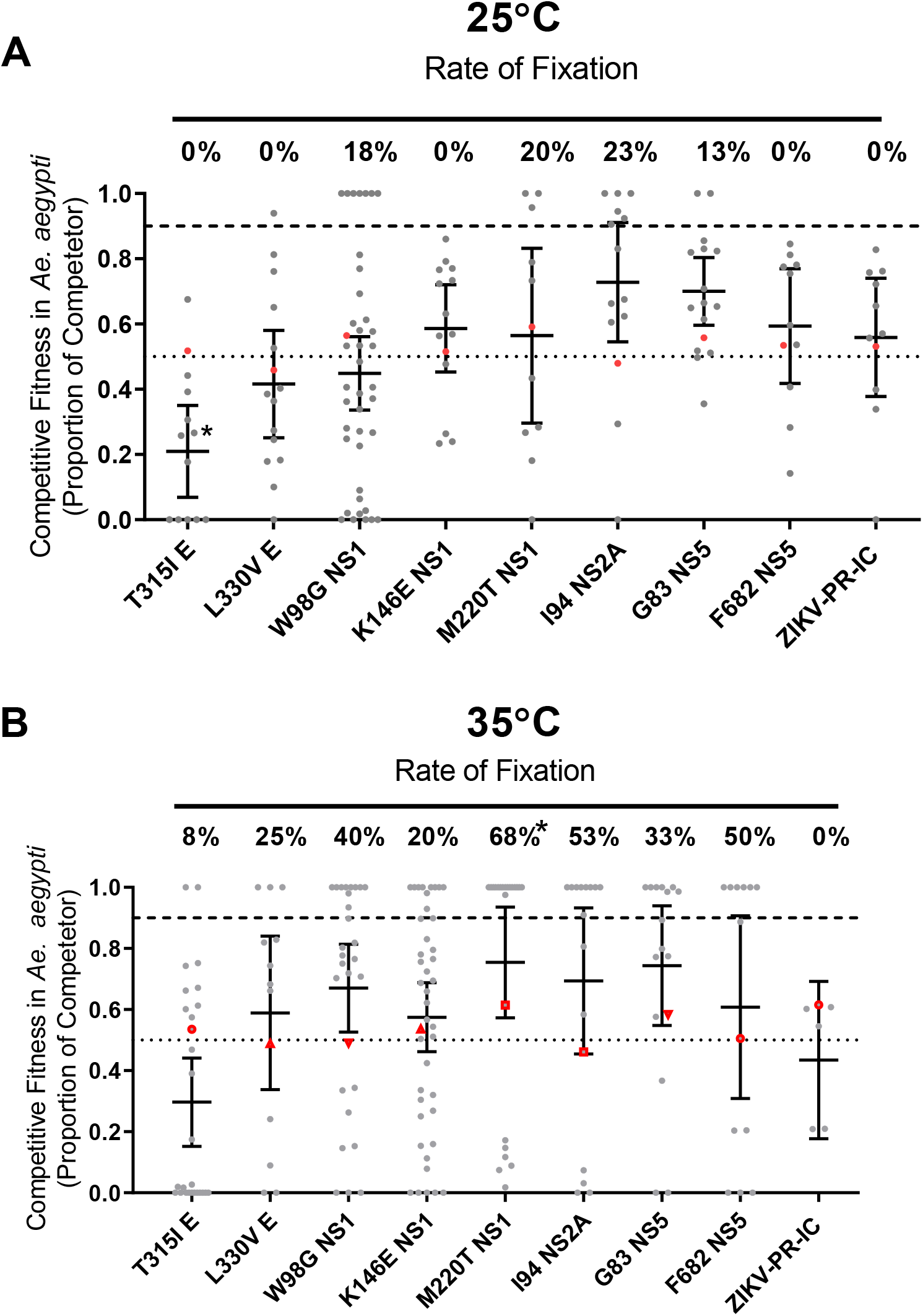
High temperature increases variant fixation in Aedes aegypti. Indicated mutations were engineered into a ZIKV-PR-IC and mixed with a ZIKV-REF virus. The proportion of each competitor (grey, mean with 95% CI, *p-value < 0.05 compared with ZIKV-PR-IC, Kruskal-Wallis and Dunn’s) and rate of fixation (*p-value < 0.05 compared with ZIKV-PR-IC, Two-tailed Fisher’s exact test) was determined from mosquito bodies at 14-dpi for *Ae. aegypti* mosquitoes held at constant EIT’s 25°C (A) & 35°C (B). Fixation indicate that 100% of the sequenced nucleotides were from the competitor virus. Initial viral inoculum (ratio of competitor virus to reference) is shown in red symbols.

Several mutants tended to rise in frequency at both 25°C and 35°C, with the overall rate of fixation (mutants that reached 100% frequency as measured by our assay) higher in mosquitoes held at 35°C (p<0.001, Two-tailed Fisher’s exact test). The fitness implications of these mutants tended to be variable. For example, in orally exposed *Ae. aegypti* bodies, the clone bearing the NS1 M220T mutant had a significant (p<0.05, Kruskal-Wallis and Dunn’s) fitness advantage 14 days after blood feeding compared to wildtype. Conversely, the envelope T315I mutant had significantly decreased (p<0.05, Kruskal-Wallis and Dunn’s) fitness compared to ZIKV-PR-IC at 25°C.

## Discussion

### Extrinsic incubation temperature alters vector competence

Vector competence is largely determined by barriers to infection and escape from mosquito midgut and salivary glands [4, 34]. Our results support the extensive existing literature [8, 9, 35] that EIT controls infection and escape mechanisms, resulting in a unimodal distribution of VC values: Extreme low 25°C and high 35°C temperatures had the lowest VC, while moderate temperatures of 28°C and 32°C had peak VC (Fig 1). These results agree with previous studies [8] and mechanistic models predicting the ZIKV thermal optimal limit of 29°C for *Ae. aegypti* [8, 36]. However, the VC of mosquitoes held at diurnal temperatures were consistently lower than the moderate temperatures of 28°C and 32°C despite having a similar mean temperature. The reasons for this are unlikely caused by direct effects of temperature on virus replication because ZIKV readily undergoes replication in vertebrates that commonly maintain temperatures of 37°C, higher than any temperature tested here. It seems more likely that the depressed VC observed at higher temps is due to indirect alterations to some aspect of the mosquito environment caused by thermal stress. While our study does not capture behavioral and physiological adaptations of mosquitoes to high temperatures, our data clearly demonstrate that rapidly fluctuating temperatures negatively influence mosquito vector competence for ZIKV compared to optimal constant temperatures.

### Temperature and vector species alter the selective environment within mosquitoes

Our data on intrahost population structure during systemic infection provides several novel insights into how temperature alters the selective environment in epidemiologically relevant arbovirus vectors. First, our data demonstrate that, in general, increasing temperature leads to increases in nucleotide diversity (Fig 2A). This may simply reflect faster virus replication at higher temperatures, with additional rounds of replication introducing additional mutations in a somewhat clocklike fashion. Alternatively, it may be that this increase in nucleotide diversity reflects the strength of positive selection at higher temperatures. The exception to this is mosquitoes that were held at 28°C. While we cannot currently explain the higher diversity in virus from these mosquitoes, it is notable that our colony has been maintained at 28°C for more than 25 generations. Mosquitoes held at diurnally fluctuating temperatures, as above, were somewhat atypical in that they tended to have higher levels of nucleotide diversity than one would expect given the distribution of diversity observed in mosquitoes held at constant temperature. As in the case of VC, diurnal temperature fluctuations are distinct in their impacts on virus-host interactions when compared to constant temperatures and thus the impact and strength of selection is also distinct.

Analysis of synonymous and nonsynonymous variation at the intrahost level revealed that signatures of strong purifying selection were observed only in mosquitoes held at diurnal temperatures (Fig 2B-D; Fig 3E, J). This observation provides an important counterpoint to several previous studies that have documented a relaxation of purifying selection in flaviviruses that are undergoing replication in mosquitoes [37]. Data from mosquitoes held at constant temperatures supports these prior findings: *d_N_/d_S_* was generally near or above 1. In some cases, (e.g. 28°C and 35°C) strong positive selection was detected within the E and NS1 coding regions. Positive selection observed at 35°C seems likely to be due to virus adaption to some component of the mosquito stress response such has heat shock protein [38, 39]. The mechanisms that give rise to the signal of positive selection at 28°C are not clear but may be due to a bottleneck that by chance resulted in a genome with one or more nonsynonymous variants that rose in frequency due to this stochastic event rather than natural selection. The strong purifying selection observed in mosquitoes held at diurnal temperatures is detectable mainly in nonstructural coding sequences, and principally in NS5. This seems logical given the role of NS5-encoded proteins in virus replication [40] and highlights the requirement for replicase functionality across a wide temperature gradient. Indeed, our data suggest that ZIKV NS5 is adapted to temperature variation rather than to any individual temperature.

Analysis of consensus sequences taken from different mosquito tissues allowed us to identify the impacts of virus dissemination and physiological barriers on ZIKV sequences during systemic infection. Generally, our data demonstrate a positive correlation between constant temperature and nucleotide diversity (Fig 2E). As the virus disseminates into saliva, nucleotide diversity increases both with temperature and in a stepwise manner as new tissue compartments are infected. These findings indicate that at higher temperatures, more new consensus variants are generated. The impact of diurnal temperatures differed between two species, with most new consensus variants generated at diurnal temperatures by the less-efficient vector *Ae. albopictus.* These observations demonstrate that temperature gradients and fluctuations, and virus transmission by new, perhaps less efficient vector species, can drive the emergence of new virus variants during mosquito infection, and are supported by analyses of divergence that incorporate both intra- and interhost variation. Therefore, migration of arboviruses such as ZIKV into new environments containing unexplored vectors represents an opportunity for the emergence of new virus variants.

### Increased extrinsic incubation temperatures drive viral variant fixation in mosquitoes

We identified 8 consensus mutations (5 non-synonymous and 3 synonymous) in multiple mosquitoes during the course of this study. These all were present in the input population at low frequencies (0.01- 0.35, Table 1) and rose in frequency during mosquito infection. Four consensus changes were found in both *Aedes* species, and in mosquitoes held under most temperature regimes. L330V E (Fig S1A) is within domain III of the envelope protein, which plays a role in host cell receptor binding and entry [41]. W98G NS1 (Fig S1B) is a surface exposed aromatic to aliphatic amino acid change on the wing section of NS1, which contributes to cellular membrane association [42]. M220T NS1 (Fig S1C) replaces a sulfur containing side group with a hydroxylic side group and is located on the loop surface the NS1 172-352 homodimer [43]. G83 NS5 (Fig S1D) is a synonymous mutation found in the middle of the coding sequence for the NS5 methyltransferase domain active site. Moreover most of these substitutions were not particularly conservative and may be of functional significance.

While none of these mutations increased significantly in frequency at 25°C, mosquito exposure to 35°C or diurnal temperatures cause some of these mutations to rise in frequency, sometimes in a species-dependent manner, again highlighting the temperature-dependence of variant frequencies in our studies.

To assess the fitness implications of these and other mutations that were repeatedly detected in mosquitoes, we engineered individual mutations into a ZIKV infectious clone and conducted *in vivo* competition studies at 25° and 35°C. The most notable finding from these studies is that higher temperatures tended to favor frequently detected mutations, which is consistent with our data on variant frequencies, consensus level changes and the strength of purifying selection. Accordingly, we conclude that the variants we examined are more likely to reach high frequency at higher temperature.

The work presented here was designed to address the hypothesis that temperature determines not only the efficiency with which mosquitoes transmit arboviruses (which has been well established for decades) but that it also influences virus evolutionary dynamics. The lack of quantitative data on how temperature effects arbovirus mutational diversity and selective forces within mosquitoes is a critical shortcoming in the literature. Data presented here provides some novel insights into this. The most significant findings reported here are related to how increases in temperature increase the rate of fixation of novel variants in virus populations, both at the population and consensus sequence levels. This suggests that as global temperatures rise, new virus variants may emerge more rapidly. This observation requires validation using other virus-vector pairs, but the implications are ominous and require further attention.

A second key finding is that diurnal temperatures impose heretofore undetected purifying selection on virus populations as they pass through mosquitoes. This finding was not predicted, but was consistent in both mosquito species examined. We suspect that this finding is related to both (a) increased constraint imposed by the requirement that the virus replicase act efficiently across a ten degree Celsius temperature range and (b) the inability of potentially temperature-specific adaptive mutations to rise in frequency. Our data on individual mutations in various mosquito tissues supports this. Moreover, this work collectively highlights the significance of temperature changes on the evolutionary biology of the mosquito-virus interaction. It also indicates that studies of arbovirus-vector interactions conducted using multiple temperatures, including diurnal temperature cycles, may capture subtle yet significant evolutionary forces that act on viruses during mosquito infection.

## Methods

### Cells and Virus

African Green Monkey kidney cells (Vero; ATCC CCL-81) were maintained at 37°C and 5% CO_2_ in Dulbecco’s modified Eagle’s medium (DMEM) supplemented with 10% fetal bovine serum (FBS) and 1% penicillin-streptomycin (Pen-Strep). Zika virus strain PRVABC59 (ZIKV-PRVABC59; GenBank # KU501215) obtained from the Center for Disease Control and Prevention branch in Fort Collins, CO was originally isolated from the sera of a patient returning from travel to Puerto Rico in December 2015. The virus was isolated on Vero cells and a 4^th^ passage frozen at −80 was used for all *in vivo* and *in vitro* experiments. ZIKV-PRVABC59 infectious clone (ZIKV-PR-IC) served as a backbone for the reverse genetic platform developed by our lab [44] to introduce all point mutations. ZIKV-REF was designed using the aforementioned reverse genetic platform, incorporating 5 synonymous mutations amino acid 108-arganie and 109-serine of the prM protein coding sequence. The ZIKV-PR-IC sequence nucleotides were changed from ZIKV-PR-IC 795-CGG TCG-800 to ZIKV-REF 795-AGA AGT-200.

### Mosquitoes

*Ae. aegypti* colonies for this study were established from individuals collected in Poza Rica, Mexico [45] and used at F13-F18 generation. A lab adapted colony (greater than 50 generations) of *Ae. albopictus* were established from individuals collected in Florida, USA; the colony was provided by the Centers for Diseases Control and Prevention (CDC-Fort Collins, CO, USA) in 2010. Mosquitoes were reared and maintained at 27-28°C and 70-80% relative humidity with a 12:12 L:D photoperiod. Water and sucrose were provided ad libitum.

### Infection of *Aedes* mosquitoes and sample collection

Adult mosquitoes used for experiments were 3-7 days post-eclosion. Mosquitoes were provided a bloodmeal containing calf blood mixed 1:1 with ZIKV-PRVABC59 (1E7 PFU/mL) using a water jacketed glass membrane feeder. Engorged female mosquitoes were sorted into cartons and housed at 25°C, 28°C, 32°C, 35°C at constant temperatures or alternating between 25°C −35°C to simulate diurnal condition, with70-80% relative humidity and 12:12 L:D photoperiod. Mosquitoes were cold anesthetized in preparation for dissociations. Mosquito midguts, legs/wings, and saliva from the first batch of mosquitoes were collected after 7- and 14-days post-feed for NGS processing. Mosquito carcass, legs/wings and saliva from the second batch of mosquitoes were collected at 3, 5, 7, 10, and 14 days post-feed for assessing systemic infecting dynamics. Tissues represent infection (midgut), dissemination (legs), and transmission (saliva). Tissues were removed using forceps cleaned with 70% ethanol between samples and were homogenized in 200 μl of mosquito diluent with a stainless-steel ball bearing using a Retsch Mixer Mill 400 at 24 Hz for 45 seconds, as previously described [46]. Saliva was collected by inserting mosquito mouthparts into capillary tubes containing mineral oil for 30 to 45 minutes. Saliva in oil was removed from the capillary tube by centrifugation into 100 μl of mosquito diluent for 5 minutes at >20,000 x g. All samples were stored at −80°C until manipulation.

### Plaque assay

ZIKV stocks and infectious bloodmeal were quantified by plaque assay on Vero cell cultures seeded in 12-well plates. Duplicate wells were infected with 0.2 ml aliquots from serial 10-fold dilutions of virus stocks and infectious blood meals in media (DMEM supplemented with 1% FBS and 1% penicillin/streptomycin), and virus was adsorbed for one hour by incubating at 37 °C in 5% CO_2_. Following incubation, the inoculum was removed, and monolayers were overlaid with tragacanth-EMEM overlay containing 1x EMEM, 5x L-glutamine, sodium bicarbonate 3.75%, tragacanth 1.2%, gentamicin (25mg/ml), and Amphotericin B 40mL/L. Cells were incubated at 37 °C in 5% CO_2_ for four days for plaque development. Cell monolayers then were stained with 1 mL of overlay containing a 20% ethanol and 0.1% crystal violet. Cells were incubated at room temperature for 30-60 minutes and then gently washed and plaques were counted. Plaque assays for 3,5,7,10 and 14 days post infection (dpi) mosquitoes were performed similar to above with the following changes, 50 ul of homogenized midgut and leg tissues or 30 ul of saliva samples were added to Vero cultures in 24-well plates (final volume of 200 ul), and plaques were observed post processing.

### Viral RNA isolation

Viral RNA was extracted from 50 μl of either cell culture supernatant, homogenized mosquito tissues, or saliva-containing solution using the Mag-Bind® Viral DNA/RNA 96 kit (Omega Bio-Tek) on the KingFisher Flex Magnetic Particle processor (Thermo Fisher Scientific). Nucleic acid extraction was performed as directed by the manufacturer and eluted in 50 μl nuclease-free water. Viral RNA was then quantified by qRT-PCR using the iTaq™ Universal Probes One-Step Kit (BIO-RAD) according to manufacturer’s protocol using a forward primer (5’-CCGCTGCCCAACACAAG-3’), reverse primer (5’-CCACTAACGTTCTTTTGCAGACAT-3’), and FAM probe (5’-AGCCTACCTTGACAAGCAGTCAGACACTCAA-3’) sequences [47].

### Generation of ZIKV mutant clones

An infectious clone for ZIKV-PRVABC59 was used to generate mutants [44]. To engineer the point mutations (Table 1) into the ZIKV genome, the corresponding single nucleic acid substitution was introduced into the ZIKV-PR-IC using *in vivo* assembly cloning methods [48]. The infectious clone plasmids were linearized by restriction endonuclease digestion, PCR purified, and ligated with T4 DNA ligase. From the assembled fragments, capped T7 RNA transcripts were generated, and the resulting RNA was electroporated into Vero cells. Infectious virus was harvested when 50-75% cytopathic effects were observed (5-8 days post transfection). Viral supernatant was clarified by centrifugation and supplemented to a final concentration of 20% fetal bovine serum and 10 mM HEPES prior to freezing and storage as single use aliquots. Titer was measured by plaque assay on Vero cells. All stocks (both wildtype and infectious clone-derived viruses) were sequenced via sanger sequencing to verify complete genome sequence.

### Competition studies

Competitive fitness was determined largely as described in previous studies [49, 50]. Competitions were conducted with orally infected *Ae. aegypti* (Poza Rica) mosquitoes. Three to seven day old mosquitoes were offered a bloodmeal containing the 1:1 mixture of viruses (ZIKV-REF and ZIKV-clone of interest) at a concentration of 1 million PFU/mL and bodies were collected 14 days post blood feed. RNA was extracted as above, and amplicons were generated via qRT-PCR using iTaq™ Universal SYBR® Green One-Step Kit (BIO-RAD) according to manufacture protocol. A locked nucleic acid (LNA) forward primer was used to ensure amplicon specificity. The forward LNA primer 5’-A+CTTGGGTTGTGTACGG-3’ and reverse primer 5’-GTTCCAAGACAACATCAACCCA-3’ were used to generate amplicons for Quantitative Sanger sequencing. Genotype fitness was analyzed using polySNP software [51] to measure the proportion of the five synonymous variants present in the ZIKV-REF sequence allowing us to compare the proportion of ZIKV-REF virus to competitor virus.

### Library preparation for next-generation sequencing

Positive controls were generated in triplicate, each generated with 1 million genome equivalents of a 100% ZIKV PRVABC59 viral stock, a mixture of 90% ZIKV PRVABC59 and 10% ZIKV PA259359 (GenBank # KX156774.2), and a mixture of 99% ZIKV PRVABC59 and 1% ZIKV PA259359. The negative control was water (no template control, or NTC). Controls and sample RNA (10ul) was prepared for NGS using the Trio RNA-Seq Library Preparation Kit (NUGEN) per manufacturer standard protocol. Final libraries were pooled by tissue type and analyzed for size distribution using the Agilent High Sensitivity D1000 Screen Tape on the Agilent Tapestation 2200, final quantification was performed using the NEBNext® Library Quant Kit for Illumina® (NEB) according to manufacturer’s protocol. 150 nt paired-end reads were generated using the Illumina HiSeq4000 at Genewiz.

### NGS processing and data analysis

NGS data were analyzed using a workflow termed “RPG (RNA virus Population Genetics) Workflow”; this workflow was generated using Snakemake [52] and workflow and related documentation can be found at https://bitbucket.org/murrieta/snakemake/src. Briefly, Read 1 and Read 2 .fastq files from paired-end Illumina HiSeq 4000 data were trimmed for Illumina adaptors and quality trimming of phred scores < 30 from the 3’ and 5’ read ends using cutadapt [53]. The reads were then mapped to the ZIKV-PRVABC59 reference sequence (GenBank # KU501215) using MOSAIK [54], similar to that previously described [55]. Picard [56], Genome Analysis Toolkit (GATK) [57], and SAMtools [58] were used for variant calling preprocessing. SNV’s and inserts and deletions (INDELS) we called using LoFreq [59] with the --call-indels command; otherwise, all settings were default. Consensus sequences were generated using the .vcf files generated above and VCFtools [60]. NTC had less than 0.02% of reads mapping to ZIKV and an average of < 8x coverage across the genome indicating little to no contamination, sequencing bleed through, or index hopping (S1 Table). Only variants in the coding sequence (nt position 108-10379), with100x coverage or greater and a cut off of 0.01 frequency were used for analysis to account for low coverage (reads per genome position) in the 3’ and 5’ untranslated regions (S1 Table, Fig S2).

Data analysis was performed using custom Python and R code integrated into the RPG Workflow. Using .vcf files generated by LoFreq and .depth files generated by GATK DepthOfCoverage command, the workflow generates .csv files that provides sequencing coverage across the CDS, Shannon entropy, richness, nucleotide diversity, *d_N_/d_S_,* and F_ST_ (compared to input population) of a specified locus. Additionally, amino acid changes, synonymous (S) and non-synonymous (NS) changes, and Shannon entropy are reported by variant positions. The same scripts are called manually outside of the RPG Workflow to perform the above analysis on specific protein coding regions or to compare divergence of populations other than the input.

### Genetic diversity

All genetic diversity calculations were incorporated into Python and R code located at https://bitbucket.org/murrieta/snakemake/src/master/scripts/. In short, richness was calculated by the sum of the intrahost SNV (iSNV) sites detected in the CDS in each population. Diversity was calculated by the sum of the iSNV frequencies per coding sequence. Complexity was calculated using Shannon entropy (*S*) which was calculated for each intrahost population (*i*) using the iSNV frequency (*p*) at each nucleotide position (*s*):

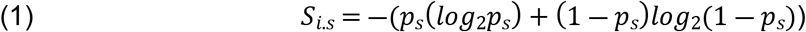

The mean S from all sites s is used to estimate the mutant spectra complexity. Divergence was calculated using F_ST_ to estimate genetic divergence between two viral populations as described previously [55]

### Selection

Intrahost selection was estimated by the ratio of nonsynonymous (*d_N_*) to synonymous (*d_S_*) SNVs per site (*d_N_/d_S_*) using the Jukes-Cantor formula as previously described [55], and incorporated into custom Python code found at https://bitbucket.org/murrieta/snakemake/src/master/scripts/. DnaSP software [61] was used to determine the number of nonsynonymous (7822.83) and synonymous (2446.17) sites from the ancestral input ZIKV consensus sequence. When no synonymous SNVs sites were present in replicates, *d_N_/d_S_* was set to 1, and no nonsynonymous SNV’s *d_N_/d_S_* was set to 0.

### Statistical analysis

All analyses were performed using GraphPad Prism (version 7.04) and R. Fisher’s exact test were used to determine significant difference in virus titers and viral loads. All other tests were done using Kruskal-Wallis with Dunn’s correction unless otherwise noted.

To evaluate the relationship between external factors and the infection dynamics of ZIKV, we examined the data with generalized linear models (Supplemental material). The predictors we used include days post infection (days), temperature (scaled), species, and tissue type. We evaluated the impact of these variables on consensus changes, vector competence, complexity, nucleotide diversity and richness. We assumed that consensus changes and richness follow a quasi-Poisson distribution, complexity and nucleotide diversity follow a linear distribution, and assumed that dissemination efficiency and vector competence follow a binomial distribution. Our original models follow the base structure

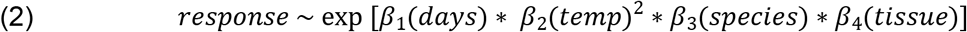

Each model was reduced to a best fit structure using AIC values and/or a chi-square goodness of fit test. The polynomial on temperature allows us to differentiate between the linear and quadratic effect of temperature. Vector competence was evaluated with the following base structure for each tissue response

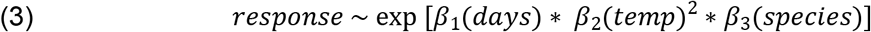

## Data Availability

Zika virus sequence data have been deposited in the NCBI Sequence Read Archive (PRJNA659260). All other data supporting the findings given are available within the article and supplementary information files, or from corresponding author upon request.

## Acknowledgments

The authors would like to acknowledge the CDC Arbovirus Reference Collection (CDC, Fort Collins) for providing the ZIKV strain used in this study. Furthermore, we would like to acknowledge Erin McDonald (CDC, Fort Collins) for providing the L330V-E and W98G-NS1 ZIKV plasmids and Irma Sanchez-Vargas (CSU) for providing the *Ae. albopictus* colony used in this study.

